# Protein Acetylation Induced by Dichloroacetate (DCA) Treatment is Associated with Decreased Respiration in Cultured Hepatocytes: Preliminary Results

**DOI:** 10.1101/662841

**Authors:** Jesse G. Meyer

## Abstract

Protein post-translational modification (PTM) by acetylation at the ε-amine on lysine residues in proteins regulates various cellular processes including transcription and metabolism. Several metabolic and genetic perturbations are known to increase acetylation of various proteins. Hyper-acetylation can also be induced using deacetylase inhibitors. While there is much interest in discovering drugs that can reverse protein acetylation, pharmacological tools that increase non-enzymatic protein acetylation are needed in order to understand the physiological role of excess protein acetylation. In this study, I assessed whether inhibition of pyruvate dehydrogenase kinase (PDHK) could cause protein hyper-acetylation due to excess production of acetyl-CoA by pyruvate dehydrogenase (PDH). Western blot of total protein from dichloroacetate (DCA) treated hepatocytes with anti-acetyl-lysine antibody showed increased protein acetylation, and seahorse respirometry of DCA pretreated hepatocytes indicated a subtle decrease in basal and maximal respiratory capacity.

## INTRODUCTION

Protein diversity from the roughly 20,000 human genes is augmented after transcription through multiple processes, including alternative splicing^1^ and protein PTMs^2^. Modern mass spectrometry technology has enabled the detection and quantification of this protein diversity^3^. In particular, significant progress has been made in detecting PTMs; mass spectrometry-based proteomic studies now routinely detect and quantify thousands of PTM sites.

One of the most well-studied groups of PTMs is lysine ε-acylation^4^. Proteins can be acylated on lysine residues by almost any small molecule that exists as a CoA conjugate, or even by other proteins such as SUMO and Ubiquitin^5^. Acylation can be either enzymatic (such as from p300/CBP^6^) or non-enzymatic. Non-enzymatic acylation results from high local concentrations of reactive carbon species^7^. Protein acylation is important because there are several examples of altered protein function resulting from acylation. For example, recent work showed that uncoupling protein 1 (UCP1) is succinylated at K56 and K151, which decreases UCP1 uncoupling activity^8^.

Several seemingly disparate conditions are reported to increase mitochondrial protein acetylation, including genetic deletion of Sirt3^9^, calorie restriction^10^, ketogenic diet^11^, chronic ethanol consumption^12^, and chronic high fructose or glucose consumption^13^. Nuclear histone acetylation is also induced by metabolism and is linked to ACLY protein activity, which makes acetyl-CoA in the cytosol from citrate^14–16^. All these conditions must increase the equilibrium level of acetylation by either altering the production of acetyl-CoA or altering the rate of lysine acetylation removal.

Protein acylation is removed by deacetylases, which include the NAD+-dependent class of deacetylates named Sirtuins^17^. Considerable effort has been devoted to the study of Sirtuin proteins since the discovery that their activity is positively correlated with lifespan^18,19^. Sirtuins isoforms exist in the nucleus^20^, cytoplasm^21,22^, and mitochondria^17^. It is now widely accepted that the various isoforms of Sirtuins remove different types of acyl groups, including acetylation, succinylation, malonylation, and palmitoylation.

Acetyl-CoA that can cause non-enzymatic protein acetylation is produced in the mitochondria by one of two paths: (1) from glycolytically-produced pyruvate by PDH, or (2) from beta oxidation of fatty acids by trifunctional enzyme. In this report, I describe preliminary results that suggest protein acetylation is increased by treatment with DCA, which increases production of mitochondrial acetyl-CoA by inhibition of PDHK^23^. PDHK inhibits PDH by phosphorylation^24,25^; DCA treatment removes the brakes that slow the entry of carbon from glycolysis into the TCA cycle. Functional analysis of DCA-pretreated hepatocytes with oxygen flux analysis suggests that DCA-induced hyperacetylation is associated with subtle decreases in basal and maximal respiration.

## METHODS

### Drug treatments

FL83B and HepG2 cells were purchased from ATCC. Cells plated in triplicate wells of 6-well plates were starved for 24 hours in base DMEM (deplete of pyruvate, glucose, and glutamine) supplemented with 5 mM glucose, 250 uM palmitate, 2 mM glutaGRO (Corning 25015CI), 10 mM HEPES, and 10% dialyzed FBS and treated with one of: (1) vehicle control (2) 5 mM dichloroacetate (DCA), (3) 5 mM 2-deoxyglucose, or (4) 80 uM etomoxir for 24 hours.

### Western blotting

Cells were then harvested with buffer consisting of 8M Urea, 100 mM Tris pH 8, 5 μM trichostatin A, and 5 mM Nicotinamide. Proteins in the cell lysates were separated using denaturing sodium dodecyl sulfate-polyacrylamide gel electrophoresis (SDS-PAGE) with 17-well Novex Tris-Glycine 4-20% Mini Gels by applying 120 volts for 1 hour. Proteins in the gel were then transferred to PVDF membranes. The membrane containing proteins was blocked for 1 hour at room temperature (RT) with 5% powdered milk in tris-buffered saline with 0.1% tween (TBST), washed 3 times with 10 mL of TBST for 5 minutes each, and then incubated with primary antibody (Cell Signalling Technologies, Acetylated-Lysine MultiMab Rabbit mAb mix #9814) according to the manufacturer’s instructions.

### Seahorse respirometry experiment

HepG2 cells were plated at 20,000 cells per well in 96-well seahorse plates and cultured for 48 hours in DMEM media containing 25 mM glucose, 10 mM HEPES, 1% AlbuMax (lipid-saturated BSA), and 5 mM carnitine with or without 5 mM dichloroacetate (DCA). To prime cells for fatty-acid utilization^26^, cells were then switched to substrated-limited base DMEM media (deplete of pyruvate, glucose, and glutamine) supplemented with 0.5 mM glucose, 1 mM GlutaGRO, 0.5mM carnitine, and 1% FBS. Forty-five minutes before the assay, cells were washed once with KHB 111 mM NaCl, 4.7 mM KCl, 1.25 mM CaCl2, 2 mM MgSO4, 1.2 mM NaH2PO4) supplemented with 2.5 mM glucose, 0.5 mM carnitine, and 5 mM HEPES, and then incubated with the same media containing 0.1% AlbuMax at 37° C in a CO_2_-free incubator. Then the oxygen consumption rate was measured at baseline and after sequential addition of 1 μM oligomycin (inhibitor of ATP synthase), 50 mM 2-deoxyglucose (inhibitor of glycolysis), 1 μM FCCP (proton gradient uncoupler), and 0.5 μM each of Rotenone and Antimycin A (inhibitors of complex I and cytochrome C reductase, respectively).

## RESULTS

Multiple cell culture media compositions were compared to determine conditions that increase total cellular protein acetylation as assessed by anti-acetyl-lysine western blot. Changes in the carbon source or nutrient composition did not have a significant influence on the relative amount of protein acetylation (data not shown). Treatment with DCA to allow unregulated production of Acetyl-CoA from glucose carbon increased protein acetylation, whereas inhibition of glycolysis with 2-DG treatment appeared to decrease protein acetylation (**Figure 1A, 1B**). Histone acetylation signal in the low molecular weight region of the membrane was mostly unchanged across samples from FL83B treated cells (**Figure 1C**), but a subtle increase in acetylated histone signal was apparent in samples from DCA- and Etomoxir-treated HepG2 cells (**Figure 1D**). Ponceau staining of the PVDF membrane showed relatively consistent loading of total protein across treatments (**Figure 1E, 1F**).

**Figure 1:**
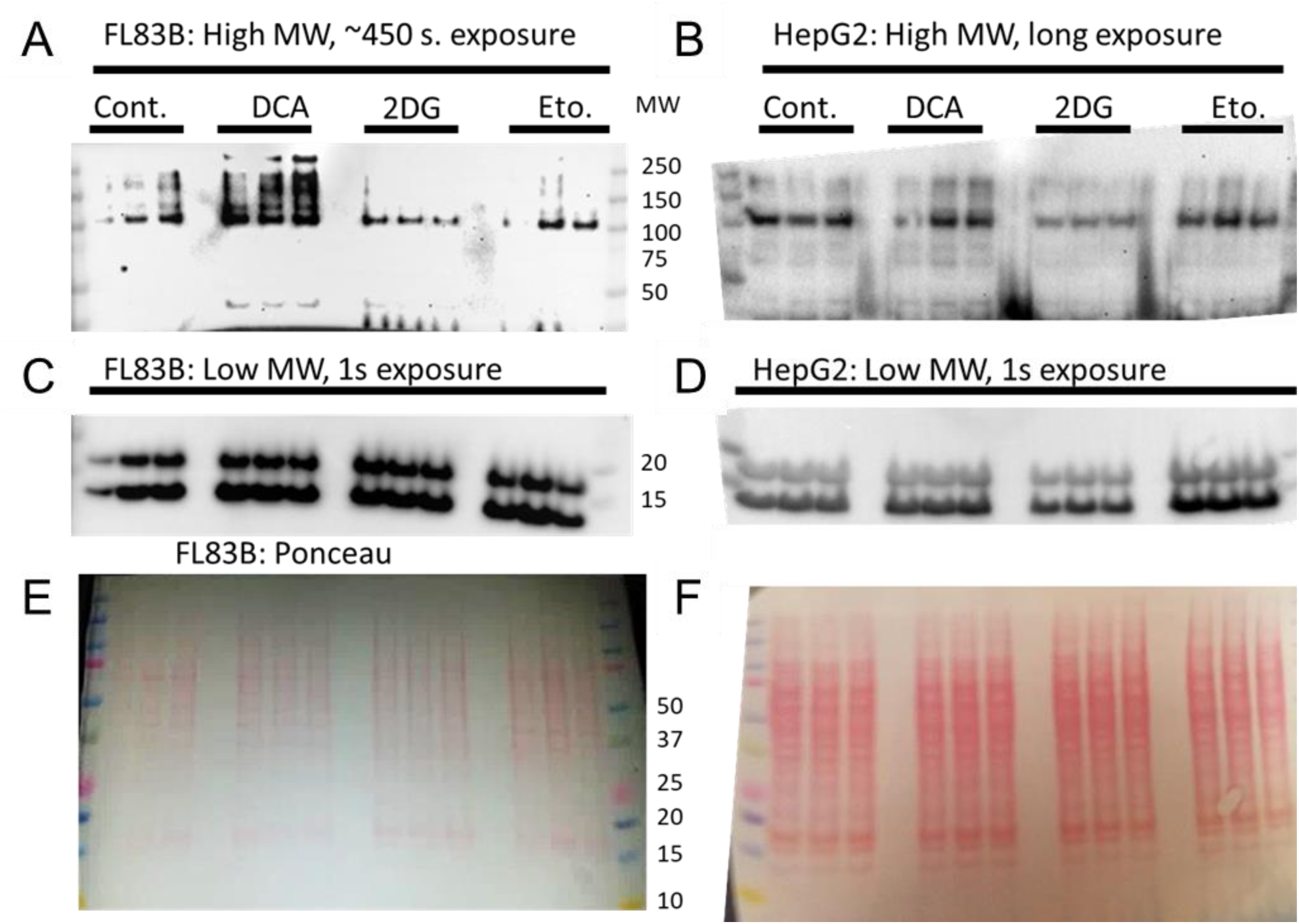
Dichloroacetate (DCA) treatment induces protein hyperacetylation in cultured hepatocytes. Triplicate samples of FL83B or HepG2 hepatocytes were treated with (1) vehicle control (2) 5 mM dichloroacetate (DCA), (3) 5 mM 2-deoxyglucose, or (4) 80 uM etomoxir for 24 hours. High molecular weight region of anti-acetyl-lysine western blot from **(A)** FL83B mouse hepatocytes and **(B)** HepG2 human hepatocytes. Low molecular weight region of the same blot membranes from **(C)** FL83B and **(D)** HepG2 cells. **(E, F)** Ponceau stain of the same membranes shown in **A-D**.

To determine whether DCA-induced protein hyperacetylation has a functional influence on the cellular metabolism, seahorse respirometry was used to measure oxygen consumption of cultured cells (**Figure 2A**). Cells were pretreated with DCA for 48 hours and then the drug was removed, and cells were exchanged into substrate-limited media to prime cells for fatty acid utilization. Oxygen consumption was measured in media containing carnitine and lipid saturated BSA. The oxygen consumption rate was corrected for total protein in each well as measured by BCA assay. The basal and maximal respiration of DCA-pretreated cells was significantly lower than controls (**Figure 2B, 2C**).

**Figure 2:**
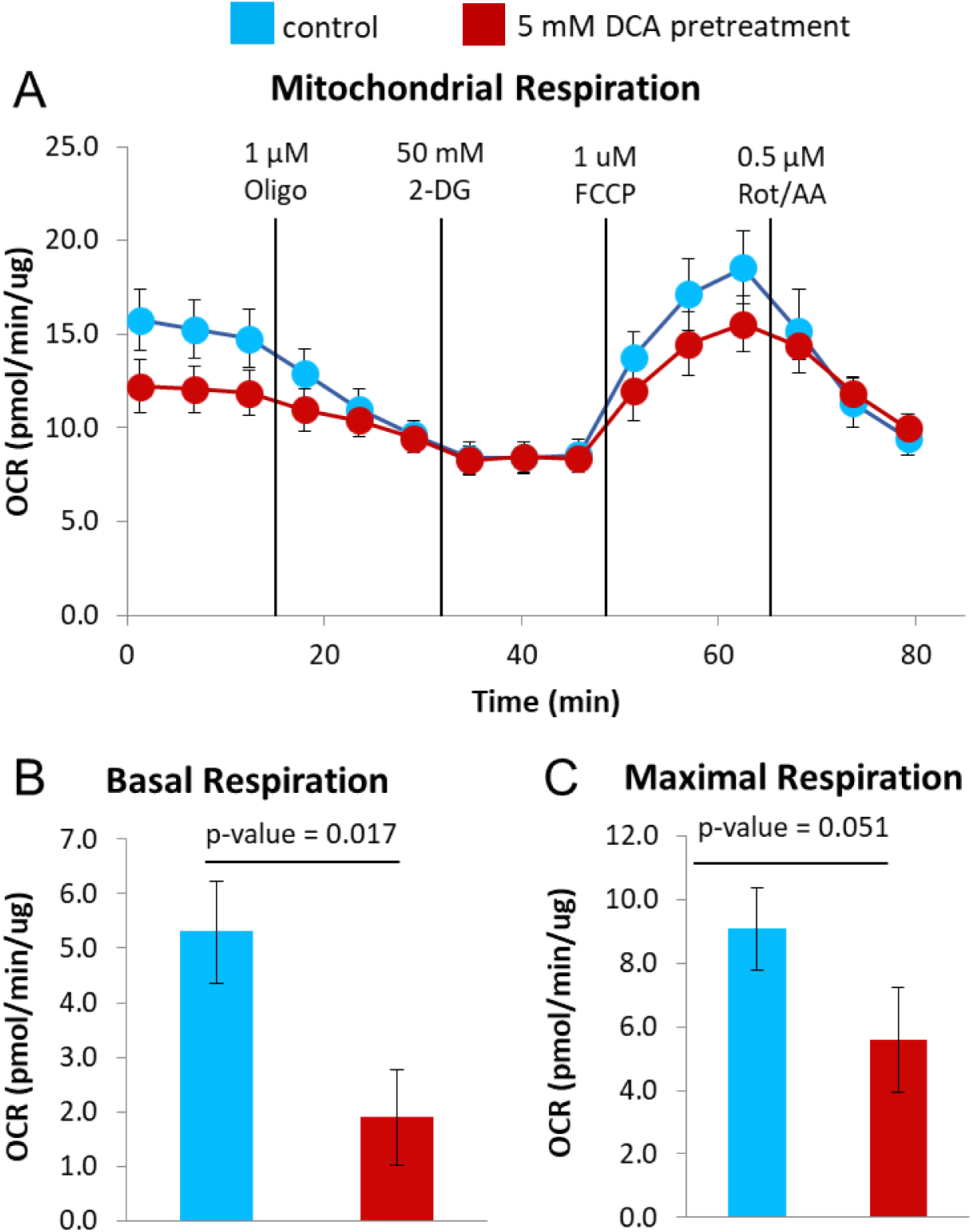
DCA-induced protein hyperacetylation is associated with decreased mitochondrial respiration in cultured HepG2 cells. **(A)** Oxygen consumption rate measured by Seahorse respirometry of HepG2 cells pretreated with control media or media containing 5 mM dichloroacetate to induce mitochondrial protein hyperacetylation. Quantification of **(B)** basal and **(C)** maximal respiration rates. n=5 wells per group, error bars show standard error measurement (SEM). p-values are from one-tailed paired t-tests.

## DISCUSSION

The results presented here suggest that unregulated flow of carbon from pyruvate into the mitochondria is sufficient to induce cellular protein hyperacetylation. Although it is likely that the hyperacetylated proteins are primarily mitochondrial due to their proximity to the production of acetyl-CoA from pyruvate, additional experiments with mass spectrometry are needed to determine the exact protein positions that are hyperacetylated. The results presented here further show that this protein hyperacetylation is associated with decreased mitochondrial respiration, which is consistent with previous reports that hyperacetylation slows mitochondrial metabolism^27^. Because media containing DCA was removed 24 hours before seahorse analysis, this altered respiration must result from cellular memory of the treatment, perhaps mediated by protein hyperacetylation.

Although the decreased respiration observed here is associated with protein hyperacetylation, there are many potentially confounding processes that may also explain this change. More work is needed to validate this effect in other cell lines and other metabolic conditions. Due to limited resources available for this study, the results reported here are preliminary, and should serve as a starting point for future studies rather than an assertion that these observations are a ground truth. Still, the data does suggest that carbon from glycolysis can be a primary driver of protein acetylation in agreement with my previous report^13^. DCA treatment may be an effective new pharmacological tool that enables deeper study of cellular protein hyperacetylation.

This preliminary work lays the groundwork for multiple future experiments. As suggested above, these experiments with DCA, 2-DG, and etomoxir treatment should be repeated with media containing ^13^C-glucose followed by mass spectrometry-based bottom-up proteomics workflows^28^ in order to verify that protein acetylation sites in proteins are indeed increased, and determine the identity of those sites that are most influenced. By using heavy glucose, this experiment would also provide definitive proof as to whether or not carbon from glycolysis can, in fact, serve as the primary carbon source for protein acetylation as suggested previously^13^. This would contradict a previous assertion that carbon for protein acetylation is exclusively from fatty acids^29^. Next, an array of media conditions with and without the same drugs should be used to treat cultured cells for oxygen flux analysis. For example, are the effects DCA-induced acetylation like those caused by Sirt3 inhibition with NAD+ and/or trichostatin A? Finally, it would be fascinated to administer DCA to mouse models and observe their respiratory exchange ratio with CLAMS cages.

## ACKNOWLEDGEMENTS

This work was funded by the National Institutes of Health NIDDK (R24 DK085610), NIA (T32 AG000266), and NLM (T15 LM007359).

## REFERENCES

1. Nilsen, T. W. & Graveley, B. R. Expansion of the eukaryotic proteome by alternative splicing. Nature 463, 457–463 (2010).

2. Olsen, J. V. & Mann, M. Status of Large-scale Analysis of Post-translational Modifications by Mass Spectrometry. Mol. Cell. Proteomics 12, 3444–3452 (2013).

3. Aebersold, R. & Mann, M. Mass-spectrometric exploration of proteome structure and function. Nature 537, 347–355 (2016).

4. Basisty, N., Meyer, J. G., Wei, L., Gibson, B. W. & Schilling, B. Simultaneous Quantification of the Acetylome and Succinylome by ‘One-Pot’ Affinity Enrichment. PROTEOMICS 1800123 (2018). doi:10.1002/pmic.201800123

5. Lumpkin, R. J. et al. Site-specific identification and quantitation of endogenous SUMO modifications under native conditions. Nat. Commun. 8, 1171 (2017).

6. Weinert, B. T. et al. Time-Resolved Analysis Reveals Rapid Dynamics and Broad Scope of the CBP/p300 Acetylome. Cell 174, 231–244.e12 (2018).

7. Trub, A. G. & Hirschey, M. D. Reactive Acyl-CoA Species Modify Proteins and Induce Carbon Stress. Trends Biochem. Sci. 43, 369–379 (2018).

8. Wang, G. et al. Regulation of UCP1 and Mitochondrial Metabolism in Brown Adipose Tissue by Reversible Succinylation. Mol. Cell 74, 844–857.e7 (2019).

9. Rardin, M. J. et al. Label-free quantitative proteomics of the lysine acetylome in mitochondria identifies substrates of SIRT3 in metabolic pathways. Proc. Natl. Acad. Sci. 110, 6601–6606 (2013).

10. Schwer, B. et al. Calorie restriction alters mitochondrial protein acetylation. Aging Cell 8, 604–606 (2009).

11. Newman, J. C. et al. Ketogenic Diet Reduces Midlife Mortality and Improves Memory in Aging Mice. Cell Metab. 26, 547–557.e8 (2017).

12. Picklo, M. J. Ethanol intoxication increases hepatic N-lysyl protein acetylation. Biochem. Biophys. Res. Commun. 376, 615–619 (2008).

13. Meyer, J. G. et al. Temporal dynamics of liver mitochondrial protein acetylation and succinylation and metabolites due to high fat diet and/or excess glucose or fructose. PLOS ONE 13, e0208973 (2018).

14. Carrer, A. et al. Impact of a High-fat Diet on Tissue Acyl-CoA and Histone Acetylation Levels. J. Biol. Chem. 292, 3312–3322 (2017).

15. McDonnell, E. et al. Lipids Reprogram Metabolism to Become a Major Carbon Source for Histone Acetylation. Cell Rep. 17, 1463–1472 (2016).

16. Wellen, K. E. et al. ATP-Citrate Lyase Links Cellular Metabolism to Histone Acetylation. Science 324, 1076–1080 (2009).

17. Carrico, C., Meyer, J. G., He, W., Gibson, B. W. & Verdin, E. The Mitochondrial Acylome Emerges: Proteomics, Regulation by Sirtuins, and Metabolic and Disease Implications. Cell Metab. 27, 497–512 (2018).

18. Kaeberlein, M., McVey, M. & Guarente, L. The SIR2/3/4 complex and SIR2 alone promote longevity in Saccharomyces cerevisiae by two different mechanisms. Genes Dev. 13, 2570–2580 (1999).

19. Guarente, L. Sir2 links chromatin silencing, metabolism, and aging. Genes Dev. 14, 1021–1026 (2000).

20. Imai, S., Armstrong, C. M., Kaeberlein, M. & Guarente, L. Transcriptional silencing and longevity protein Sir2 is an NAD-dependent histone deacetylase. Nature 403, 795–800 (2000).

21. Nishida, Y. et al. SIRT5 Regulates both Cytosolic and Mitochondrial Protein Malonylation with Glycolysis as a Major Target. Mol. Cell 59, 321–332 (2015).

22. Anderson, K. A. et al. SIRT4 Is a Lysine Deacylase that Controls Leucine Metabolism and Insulin Secretion. Cell Metab. 25, 838–855.e15 (2017).

23. Whitehouse, S., Cooper, R. H. & Randle, P. J. Mechanism of activation of pyruvate dehydrogenase by dichloroacetate and other halogenated carboxylic acids. Biochem. J. 141, 761–774 (1974).

24. Kolobova, E., Tuganova, A., Boulatnikov, I. & Popov, K. M. Regulation of pyruvate dehydrogenase activity through phosphorylation at multiple sites. Biochem. J. 358, 69–77 (2001).

25. Korotchkina, L. G. & Patel, M. S. Site Specificity of Four Pyruvate Dehydrogenase Kinase Isoenzymes toward the Three Phosphorylation Sites of Human Pyruvate Dehydrogenase. J. Biol. Chem. 276, 37223–37229 (2001).

26. Wang, D., Green, M. F., McDonnell, E. & Hirschey, M. D. Oxygen Flux Analysis to Understand the Biological Function of Sirtuins. in Sirtuins (ed. Hirschey, M. D.) 1077, 241–258 (Humana Press, 2013).

27. Hirschey, M. D. et al. SIRT3 regulates mitochondrial fatty-acid oxidation by reversible enzyme deacetylation. Nature 464, 121–125 (2010).

28. Schilling, B., Meyer, J. G., Wei, L., Ott, M. & Verdin, E. High-Resolution Mass Spectrometry to Identify and Quantify Acetylation Protein Targets. in Psychotherapie 3–16 (Springer Berlin Heidelberg, 2019). doi:10.1007/978-1-4939-9434-2_1

29. Pougovkina, O. et al. Mitochondrial protein acetylation is driven by acetyl-CoA from fatty acid oxidation. Hum. Mol. Genet. 23, 3513–3522 (2014).

